# An Experiment in Learning the Language of Sequence Motifs: Sequence Logos vs. Finite-State Machines

**DOI:** 10.1101/143024

**Authors:** Alexandre P Francisco, Travis Gagie, Dominik Kempa, Leena Salmela, Sophie Sverdlov, Jarkko Toivonen, Esko Ukkonen

## Abstract

Position weight matrices (PWMs) are the standard way to model binding site affinities in bioinformatics. However, they assume that symbol occurrences are position independent and, hence, they do not take into account symbols co-occurrence at different sequence positions. To address this problem, we propose to construct finite-state machines (FSMs) instead. A modified version of the Evidence-Driven State Merging (EDSM) heuristic is used to reduce the number of states as FSMs grow too quickly as a function of the number of sequences to reveal any useful structure. We tested our approach on sequence data for the transcription factor HNF4 and found out that the constructed FSMs provide small representations and an intuitive visualization. Furthermore, the FSM was better than PWMs at discriminating the positive and negative sequences in our data set.

## 1 Introduction

High-throughput experiments characterizing transcription factor (TF) binding sites produce large sets of sequences. The standard way to model the binding affinities based on such data sets is to build a position weight matrix (PWM) [13] which assigns a weight for each nucleotide at each position. PWMs are often visualized as sequence logos [12]. One weakness of PWMs is that they cannot model dependencies between the positions. Other representations have been proposed but none have gained widespread acceptance.

We propose to use finite state machines (FSMs) to model the binding affinities and give a method for constructing them using a modified version of Evidence-Driven State Merging (EDSM) [10]. We tested the method on data for one transcription factor. The FSM built by our method was compact and thus useful for visualizing the data. Furthermore, the FSM discriminated the positive and negative sequences in the data set better than PWMs.

## 2 Data

The HT-SELEX (High-Throughput Systematic Evolution of Ligands by Exponential Enrichment) [5] starts with a (uniformly) random library of 40-mers in a solution. The DNA fragments were allowed to bind TFs and the unbound DNA was washed out. The remaining DNA was amplified and then sequenced. The process was repeated multiple times. In later cycles the binding sites of the TF have been strongly enriched in the sequences. We used the data from cycle number 3.

Throughout this paper we will use as test data HT-SELEX data for transcription factor HNF4 (Accession number: ERR194356) [6]. It contained 655 432 sequences each of length 40 bp. From this data set the most common 13-mer (GGGTCAAAGTCCA) was chosen as a seed. Then all the 13-mers that are within Hamming distance 6 from the seed and contained the sequence AAA in the middle were selected. This resulted in 54236 distinct fragments, of which 60 occurred at least 1000 times each and 34566 occurred only once each. The fragments occurring at least 1000 times form the positive set and the fragments occurring only once form the negative set in our experiments.

## 3 Position Weight Matrices

We first aligned the sequences in the positive set and built a PWM for them weighing each sequence by its frequency. We also tried extending the positive set to 200 sequences occurring most often and similarly building a PWM for this set. The PWMs are visualized as sequence logos in Figure 1. Sequence alignments were obtained with Clustal W [7]. PWMs and sequence logos were built with the MEME SUITE [8].

**Fig. 1.**
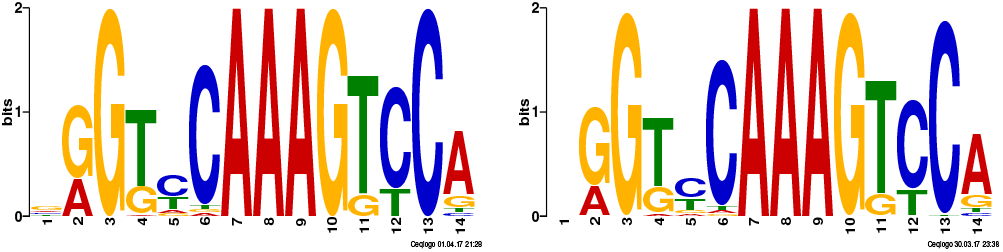
The sequence logo of the 60 DNA fragments occurring at least 1000 times each, aligned and weighted by their frequencies (**left**); the logo for the 200 fragments occurring most frequently, aligned and weighted (**right**);

The PWMs revealed some structure, such as the importance of there being three As roughly in the middle of the bound fragments, but we suspected they might mask dependencies: e.g., it might be that a poor match on one side of those As could be compensated for by a good match on the other side, but poor matches on both sides would prevent binding.

To check objectively whether fragments’ similarities to the PWMs could be used to distinguish those in the positive and negative sets, we computed the alignment cost of each fragment against each PWM. Define *false negatives* and *false positives* for a PWM with respect to a threshold to be, respectively, fragments occurring at least 1000 times each but whose alignment cost is above that threshold, and fragments occurring only once whose alignment cost is at most equal to that threshold.

Note that, since PWMs have length 14, we could not compare them directly with the fragments, which have length 13. Hence, we generated PWMs for each fragment and we used TomTom [9], also included in the MEME SUITE, to compare each PWM built for the aligned sequences against all PWMs built for the fragments. TomTom searches a database of motifs with a given query motif, considering different relative orientations and offsets, and using a method similar to the one used to match a motif against a given sequence. We excluded different orientations in our study since we know them a priori.

We computed the number of false positives required for each PWM to achieve each number of false negatives between 0 and 60; the results are shown in Table 1. We regard all the PWMs as unsatisfactory filters since, in order to accept all the positive fragments, we must also accept hundreds or thousands of negative fragments.

**Table 1.**
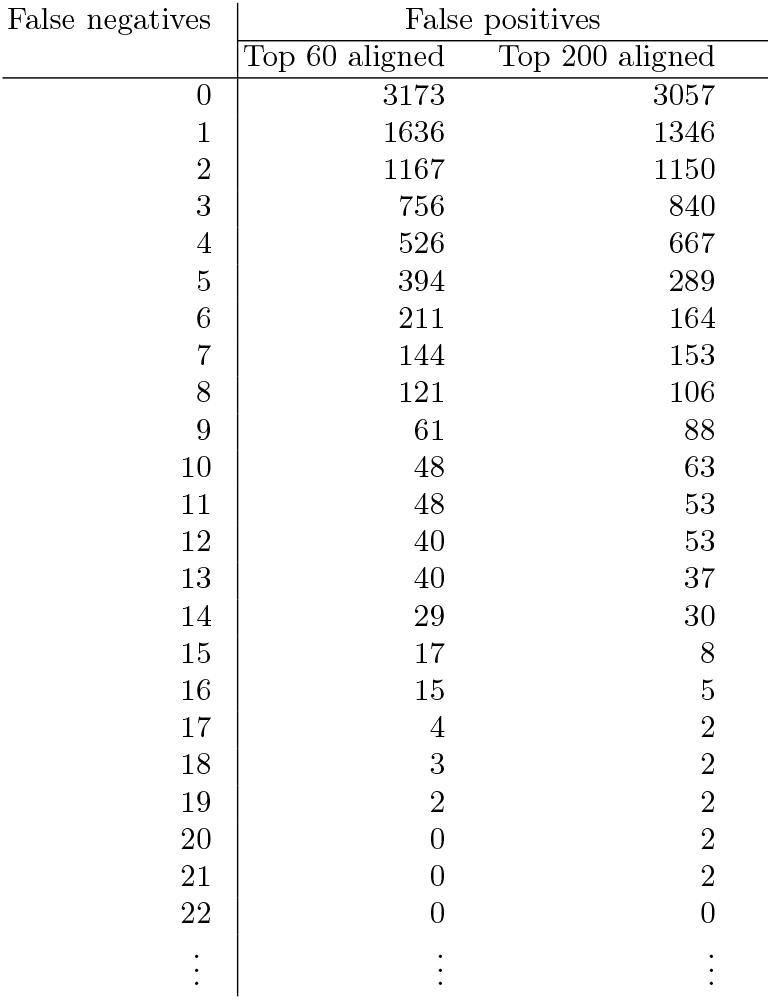
We can choose the minimum threshold such that only a given number of the 60 DNA fragments occurring at least 1000 times each, have alignment costs against a particular PWM above that threshold (false negatives). A number of the 34566 fragments occurring only once will have alignment costs less than or equal to that threshold (false positives). The number of false positives in a non-decreasing function of the number of false negatives, so all the lines after 22 contain only 0s.

## 4 Finite-State Machines

The standard way for computer scientists to represent sequential dependencies is with finite-state machines (FSMs) or high-order Markov models. We built FSMs that accepted only the most frequent fragments, but these FSMs grew too quickly as a function of the number of fragments to be able to reveal any useful structure. We therefore turned to algorithms for inferring FSMs from sets of positive and negative examples. This is a well-studied NP-hard problem [3, 1] (see, e.g., [4, 11] for recent discussions) for which there are several practical heuristics. We implemented a version of Evidence-Driven State Merging (EDSM) [10], modified such that it tries to merge only states at the same distance from the initial state, so the resulting FSM should be easier to understand.

We used as positive examples the 60 fragments each occurring at least 1000 times, and as negative examples a sample of the fragments occurring only once, each chosen with probability 0.1 thus producing about 3400 negative examples. We used fragments occurring only once because we do not know what fragments were not present in the experiment. First, we aligned the positive examples and inserted gaps in negative examples in all possible ways, so all the examples had length 14. EDSM works in rounds, starting with a trie of all the example fragments, consisting of about 20000 states, with the leaf nodes labelled accept or reject. With our version, in each round we try to merge all pairs of states at the same distance from the initial state: merging two nodes causes a cascade of implied merges, which fails if it ever requires merging an accept state with a reject state; if the whole cascade succeeds, we assign the merge an evidence score equal to the decrease in the size of the FSM; we then undo the cascade and the merge. We choose the merge with the highest evidence score, perform it, and proceed to the next round, stopping when no merge has a positive score. The size of the resulting FSM depends on the set of negative examples chosen and thus we ran the algorithm several rounds. It took several minutes to build each FSM. Still, within an hour we found an FSM with 34 states, which had four edges labelled with the gap character ‘-’, corresponding to ∊-transitions. We adjusted the FSM to remove these edges, resulting in the machine shown in the center in Figure 2: the gap-edge between states 0 and 2 we could remove by making state 2 initial; the gap-edge between states 11 and 13 we could remove by adding an edge from 11 to 15; and we could remove the gap-edges from states 30 and 31 to state 32 by making the former final. This non-deterministic 34-state FSM accepts all 60 positive examples and 57 other fragments in our initial data set, shown on the right of Figure 2, including only one fragment in the negative set. That is, the FSM is arguably better at discriminating the positive and negative sequences than a PWM, in addition to seeming subjectively more informative.

**Fig. 2.**
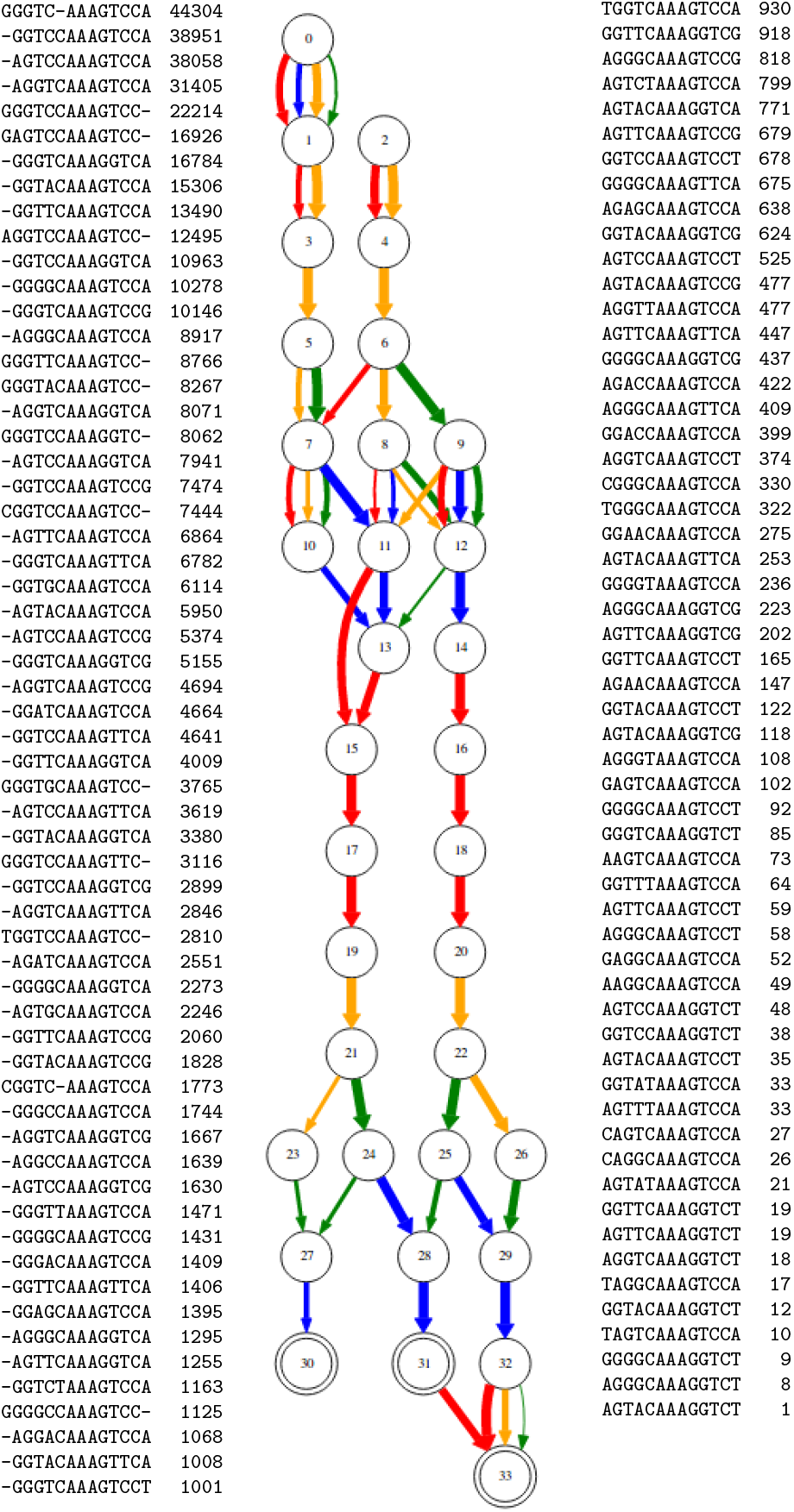
The 60 DNA fragments occurring at least 1000 times each, aligned (**left**); a 34-state non-deterministic FSM with states 0 and 2 initial, accepting all the positive examples and 57 other fragments, including one occurring only once (**center**); those 57 other fragments (**right**). In the FSMs, edges’ widths indicate how often they are crossed in accepting computations (although this does not affect the FSM’s behaviours) and their colours indicate their labels: red for **A**, blue for **C**, yellow for **G** and green for **T**.

## 5 Conclusion

We believe our investigation shows that grammatical inference, in particular building FSMs with EDSM, can produce useful models of sequence motifs and can be competitive with PWMs. Grammatical inference has been used in bioinformatics (see, e.g., [2]) but we know of no previous work quite like this. We plan to continue our studies, tuning our modification of EDSM and testing it on our current dataset and others.

## References

1 Angluin, D.: On the complexity of minimum inference of regular sets. Information and Control 39(3), 337–350 (1978)

2 Coste, F.: Learning the language of biological sequences. In: Heinz, J., Sempere, J.M. (eds.) Topics in Grammatical Inference, pp. 215–247. Springer (2016)

3 Gold, E.M.: Complexity of automaton identification from given data. Information and control 37(3), 302–320 (1978)

4 Gruber, H., Holzer, M., Jakobi, S.: More on deterministic and nondeterministic finite cover automata. Theoretical Computer Science (to appear)

5 Jolma, A., Kivioja, T., Toivonen, J. et al: Multiplexed massively parallel SELEX for characterization of human transcription factor binding specificities. Genome Research 20(6):861–873 (2010)

6 Jolma, A., Yan, J., Whitington, T., Toivonen, J. et al: DNA-Binding Specificities of Human Transcription Factors. Cell 152(1–2):327–339 (2013)

7 Larkin, M.A., Blackshields, G., Brown, N.P., Chenna, R., McGettigan, P.A., McWilliam, H., Valentin, F., Wallace, I.M., Wilm, A., Lopez, R., Thompson, J.D.: Clustal W and Clustal X version 2.0. Bioinformatics, 23(21):2947–2948 (2007).

8 Bailey, T.L., Boden, M., Buske, F.A., Frith, M., Grant, C.E., Clementi, L., Ren, J., Li, W.W., Noble, W.S.: MEME SUITE: tools for motif discovery and searching. Nucleic Acids Research, 37:W202–W208 (2009).

9 Gupta, S., Stamatoyannopoulos, J.A., Bailey, T.L., Noble, W.S.: Quantifying similarity between motifs. Genome Biology, 8(2):R24 (2007).

10 Lang, K.J., Pearlmutter, B.A., Price, R.A.: Results of the Abbadingo One DFA learning competition and a new evidence-driven state merging algorithm. In: Proceedings of the 4th International Colloquium on Grammatical Inference (ICGI). pp. 1–12. Springer (1998)

11 Vázquez de Parga, M., García, P., López, D.: A sufficient condition to polynomially compute a minimum separating DFA. Information Sciences 370, 204–220 (2016)

12 Schneider, T.D., Stephens, R.M.: Sequence logos: a new way to display consensus sequences. Nucleic Acids Research 18(20), 6097–6100 (1990)

13 Stormo, G.D., Schneider, T.D., Gold, L., Ehrenfeucht, A.: Use of the ’Perceptron’ algorithm to distinguish translational initiation sites in E. coli. Nucleic Acids Research 10(9), 2997–3011 (1982)

